## Supplemental Figures and Tables for "Protein allocation and utilization in the versatile chemolithoautotroph *Cupriavidus necator*"

for the manuscript with the title

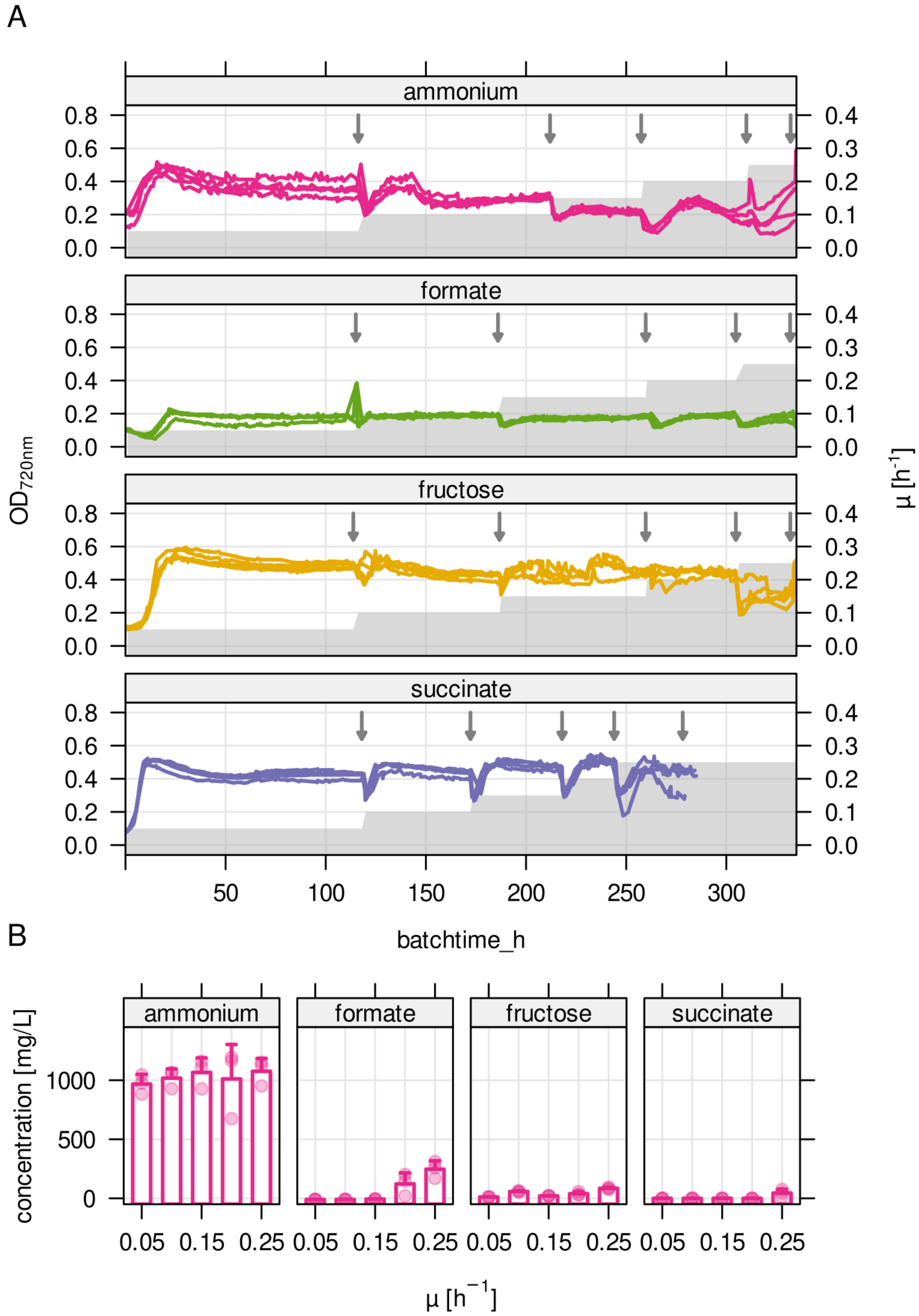


**Figure S1**. **A)** Stepwise increase in growth rate over cultivation time (grey area) and optical density at 720 nm ($OD_{720nm}$) during chemostat cultivation (4 replicates, colored lines). Arrows mark sampling time points. **B)** Residual substrate concentration for the four limiting conditions. For ammonium limitation, residual fructose concentration is shown.

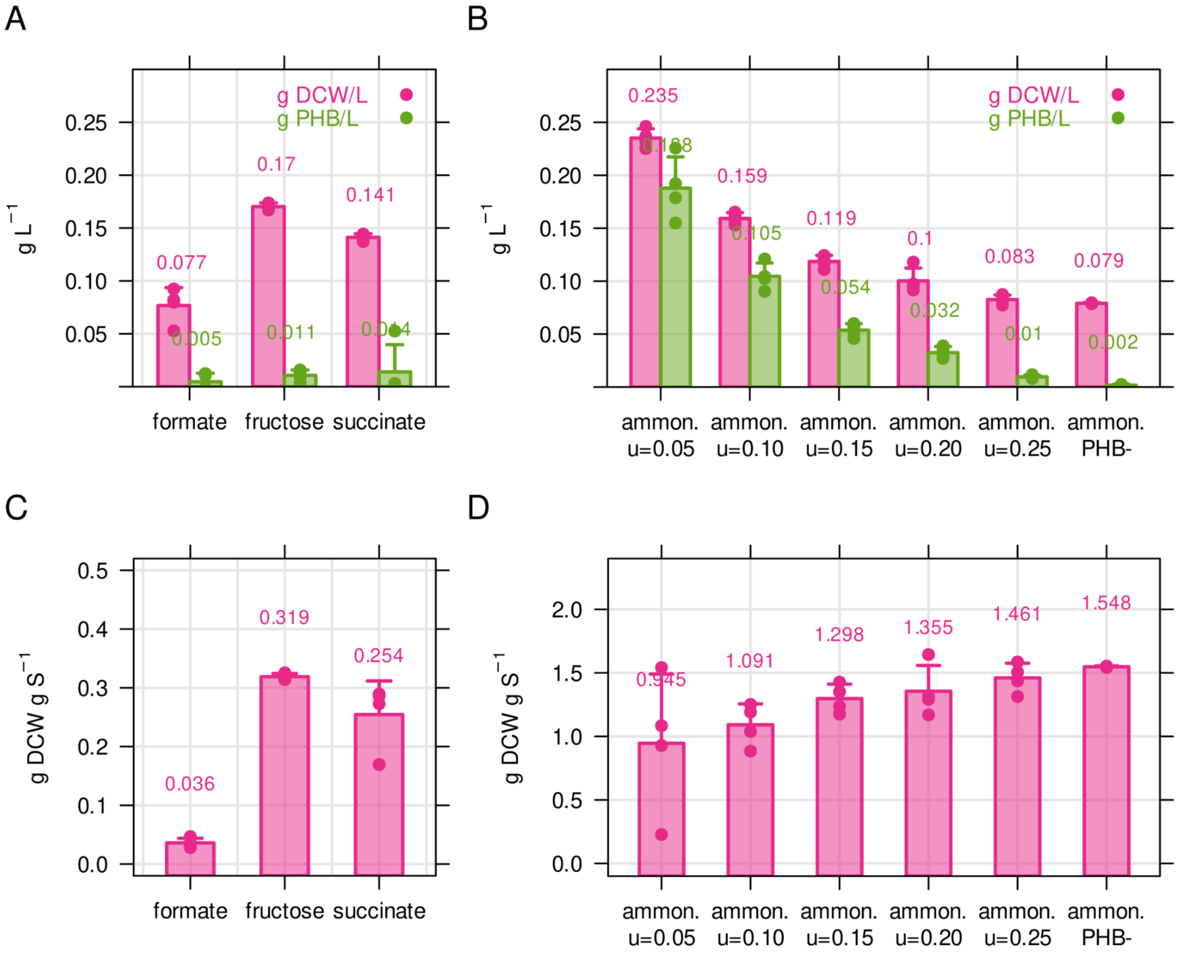


**Figure S2**. **A)** Biomass concentration and PHB content for *C. necator* in g $L^{-1}$. Cultivations were performed in 50 mL shake flasks with 2 g $L^{-1}$ formate, 0.5 g $L^{-1}$ fructose or 0.5 g $L^{-1}$ succinate. Biomass concentration was determined by dry cell weight measurement (DCW), and PHB content was determined by solvent extraction and acid hydrolysis, see Methods. **B)** Biomass concentration and PHB content for ammonium limited *C. necator* (2 g/L fructose, 0.05 g/L $NH_4Cl$). Bacteria were cultivated in chemostat bioreactors with growth rate increasing from 0.05 to 0.25 $h^{-1}$. PHB-, PHB knockout strain used as negative control. **C)** PHB-free biomass yield for three different carbon sources, calculated from A). **D)** PHB-free biomass yield for ammonium calculated from B). Bars and error bars represent mean and standard deviation of four biological replicates.

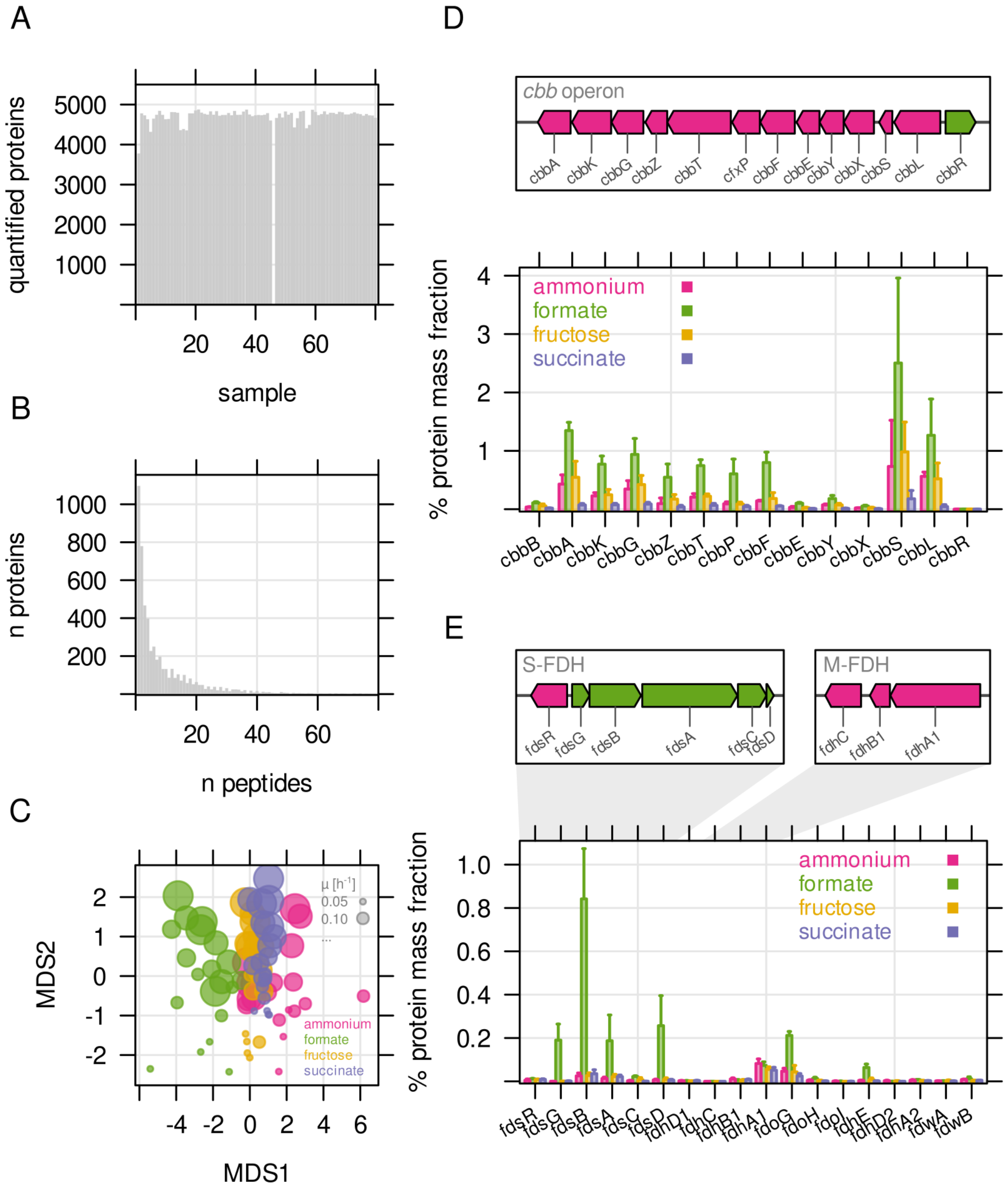


**Figure S3**. **A)** Number of quantified proteins for all 80 samples analyzed by LC-MS. Quantification for one sample failed. **B)** Number of proteins quantified by number of unique peptides. **C)** Sample similarity by protein abundance visualized using non-metric multidimensional scaling (NMDS). Each point represents one replicate sample (n = 4). Bubble size - growth rate; bubble color - type of limitation. **D)** Genome organisation of the *cbb* operon (chromosome 2) and mass fraction (%) of *cbb* proteins. The two *cbb* copies (chromosomal, pHG1) can not be distinguished by LC-MS. Protein mass fractions were therefore summed up for this representation. **E)** Protein mass fraction (%) for formate dehydrogenases. Genome maps indicate the main soluble (S-FDH) and membrane-bound formate dehydrogenase (M-FDH) operons, *fds* and *fdh*, respectively.

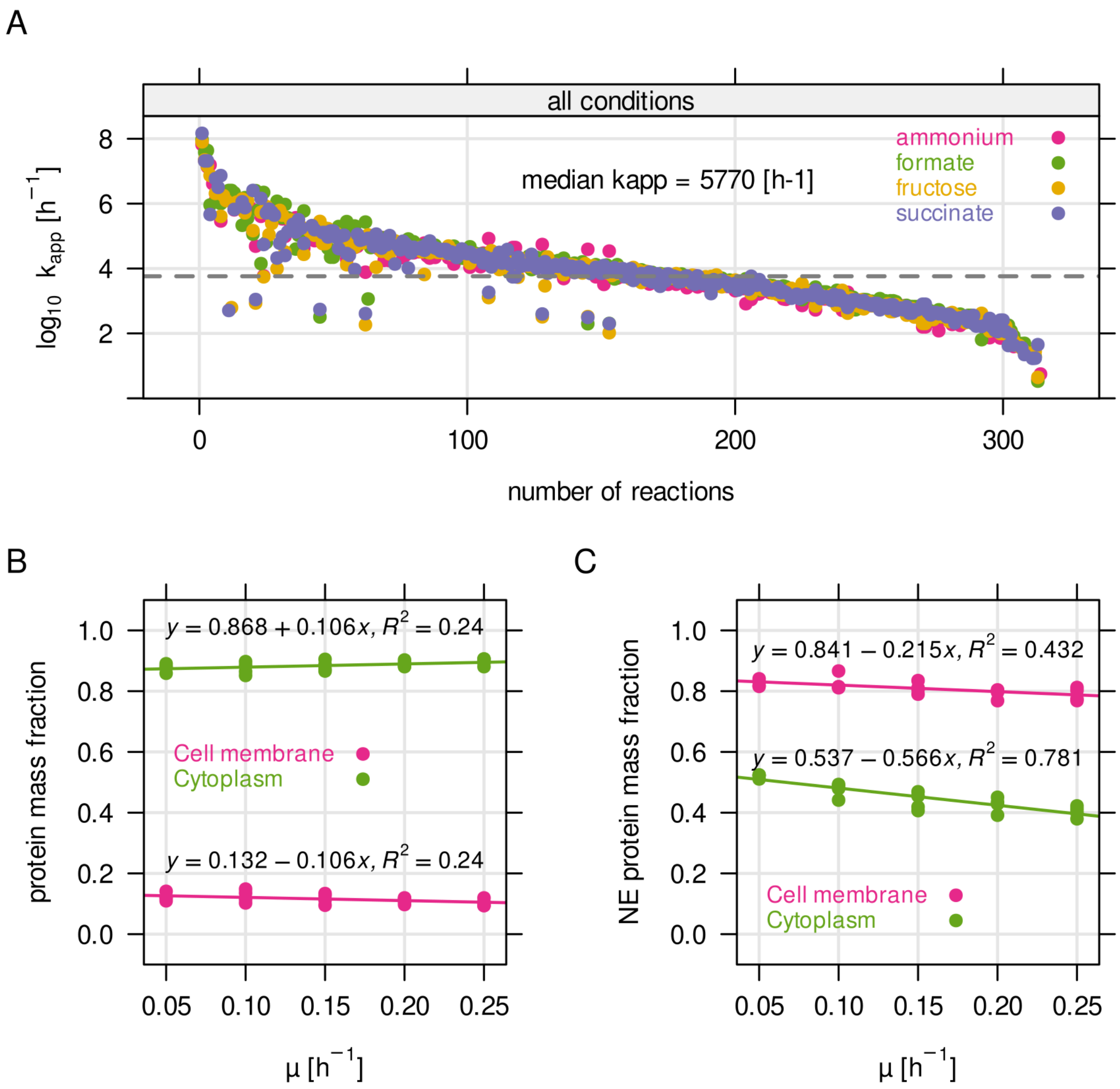


**Figure S4**. **A)** Distribution of $k_{app}$, the enzyme efficiency, obtained from parameter estimation using RBApy (Bulovic et al., 2019). Briefly, $k_{app}$ was determined by dividing the expected flux of a reaction in mmol gDCW$^{-1}$ h$^{-1}$ by the steady state enzyme abundance in g gDCW$^{-1}$ associated with the reaction. **B)** Protein mass fraction allocated to the cytoplasmic membrane and to the cytoplasm. **C)** Non-enzyme (NE) protein mass fraction, per compartment.

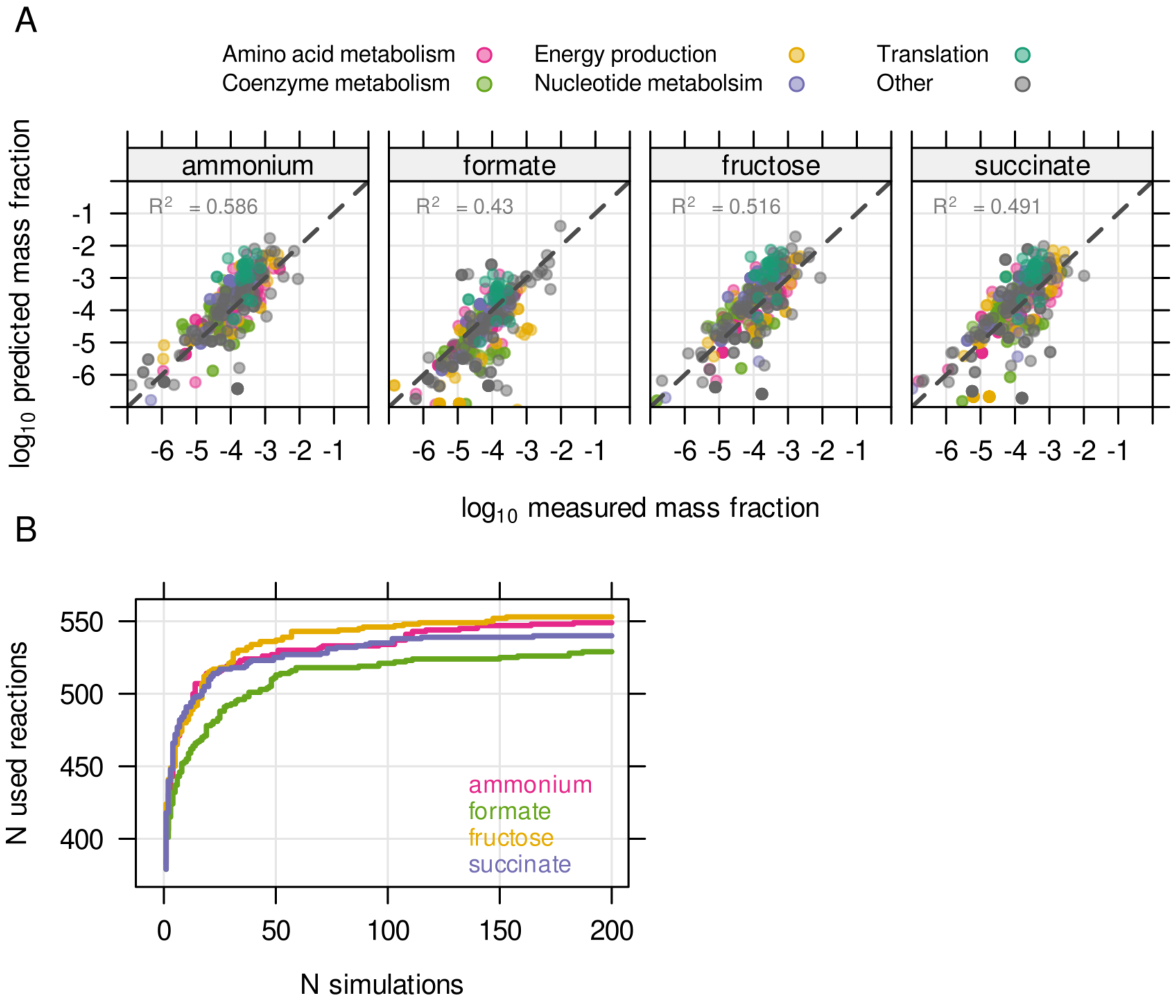


**Figure S5**. The unutilized proteome of Cupriavidus is related to environmental readiness. **A)** The resource balance analysis (RBA) model was constrained by enzyme efficiency ($k_{app}$) estimated from known protein concentrations. The model was able to reproduce measured protein allocation for four different substrate limitations. **B)** Enzyme efficiency $k_{app}$ was randomly sampled 200 times per condition to obtain the maximum number of potentially utilized reactions in each growth condition.

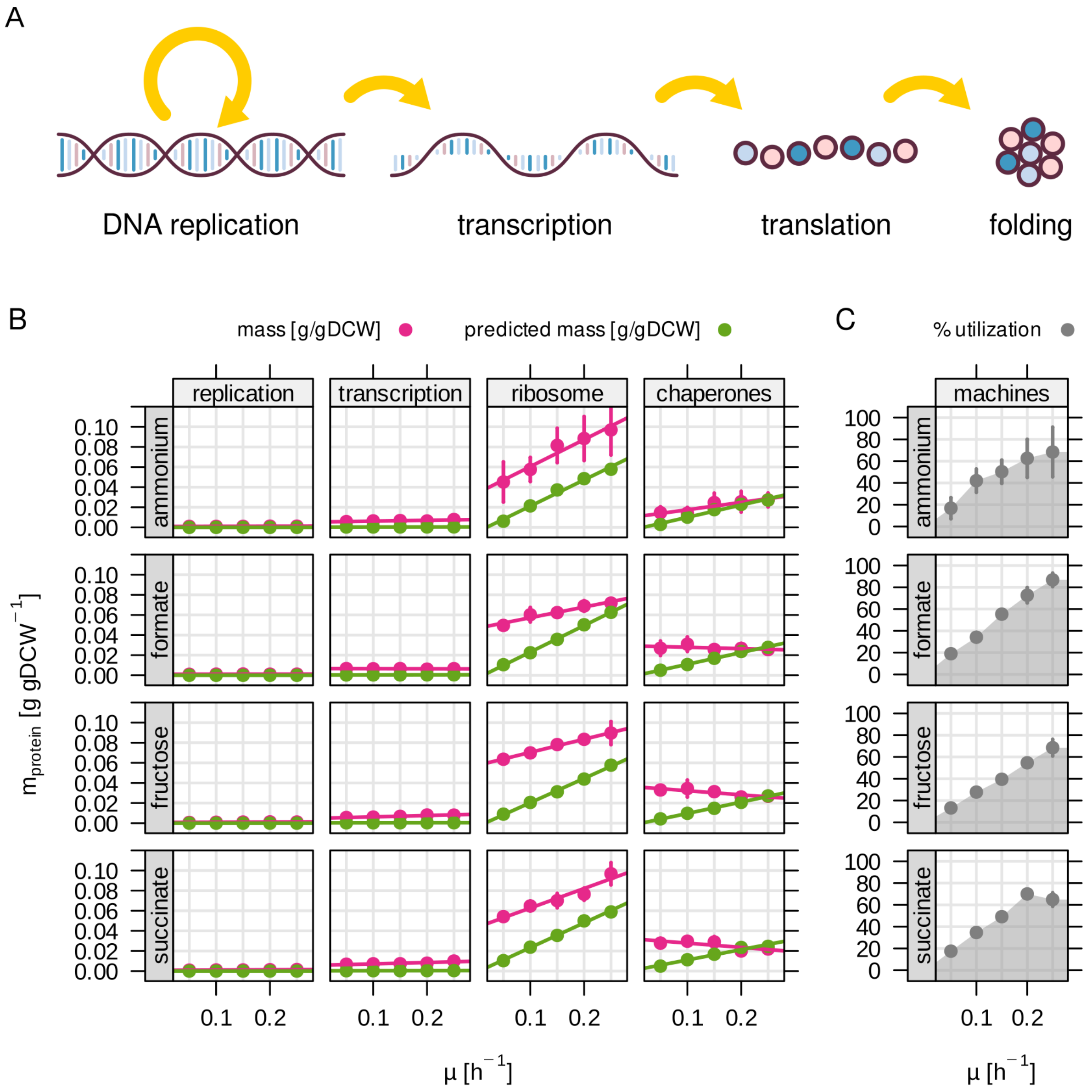


**Figure S6**. Utilization of molecular machinery related to central dogma depends on growth rate. **A)** The RBA model includes four different 'machines' for information processing. **B)** Number of unique protein subunits and associated rate for one holo-enzyme. **C)** Enzyme abundance in g/gDCW for each machine as a function of growth rate. Green - RBA model prediction, red - experimentally determined enzyme abundance. **D)** Machine utilization in % for all machine abundances summed up.

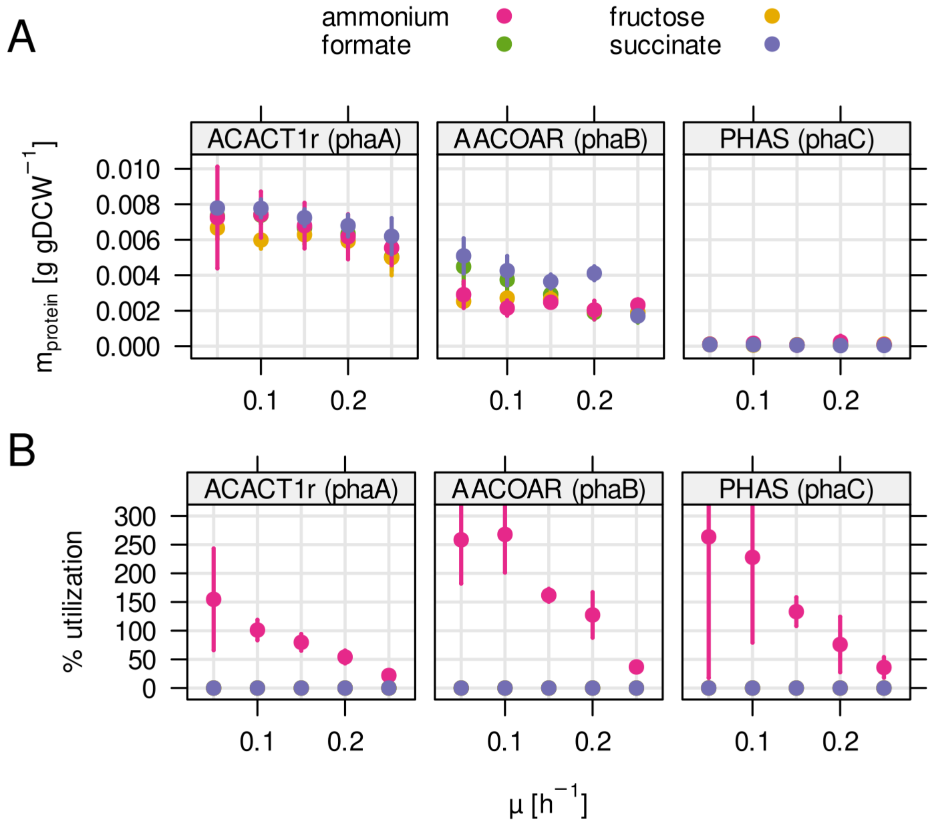


**Figure S7**. Enzyme mass and utilization of the PHB biosynthesis pathway. **A)** Protein mass fraction allocated to each enzyme during one of four different substrate limitations (ammonium, fructose, formate, succinate). **B)** Predicted enzyme utilization by RBA model simulations. Contrary to most enzymes in central metabolism, utilization increased with decreasing growth rate and was only predicted for ammonium limitation.

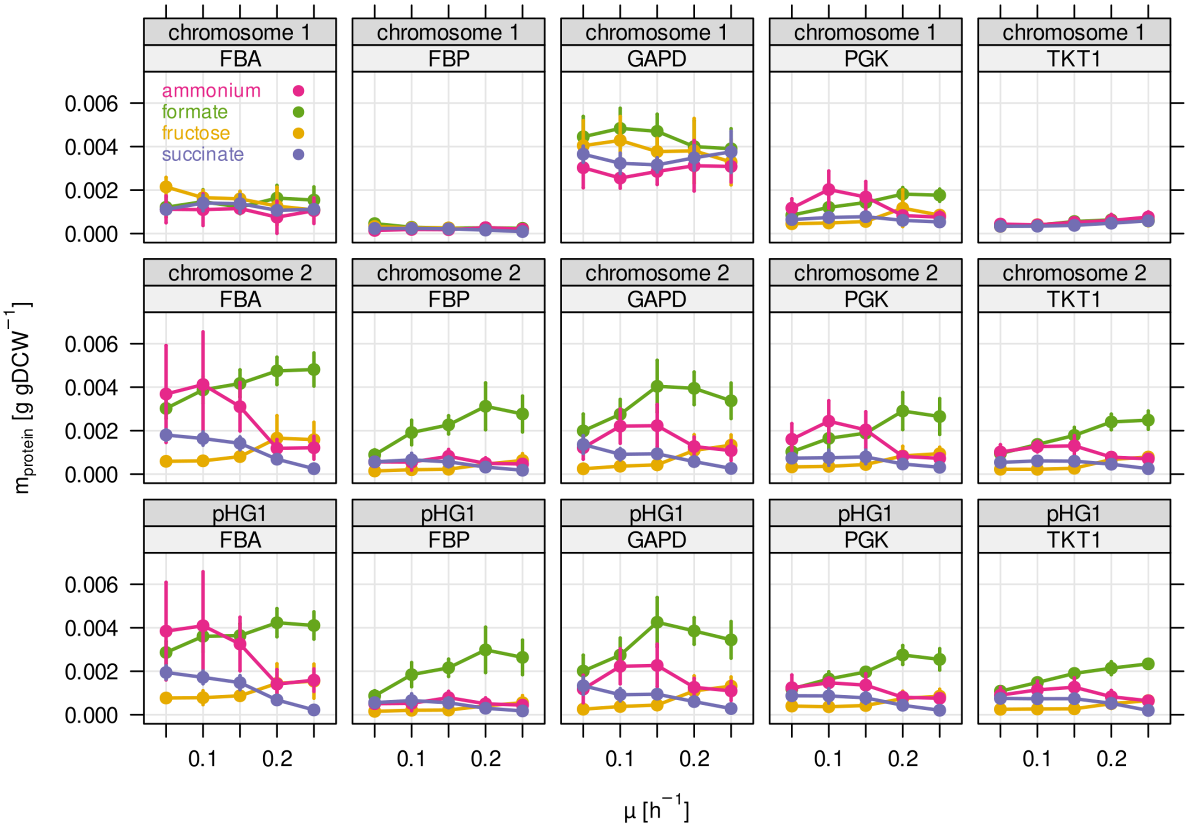


**Figure S8**. Protein mass of the five most abundant glycolytic enzymes broken down by genetic locus. Several enzymes of glycolysis/CBB cycle are present in three different copies in the *C. necator* genome. Two of these copies are located in two almost identical *cbb* operons on chromosome 2, and megaplasmid pHG1. The remaining copy on chromosome 1 is presumably the phylogenetically most ancestral, while the other copies on chromosome 2 and pHG1 were acquired later.

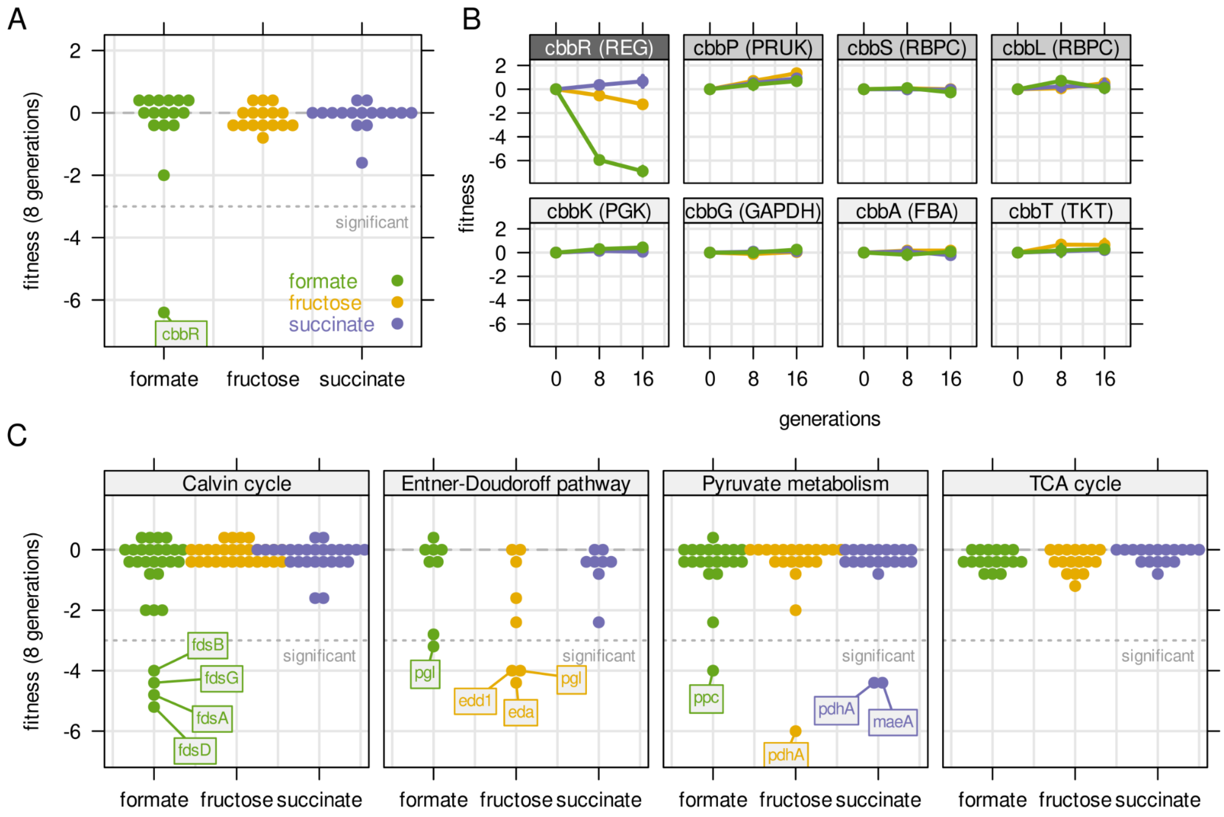


**Figure S9**. Fitness of selected genes obtained from cultivation of a barcoded transposon mutant library. The library was cultivated for 16 generations with a pulsed growth regime (dilution at 2 h intervals). **A)** Fitness for all *cbb* genes during growth on three different carbon sources. Compare also Figure 5 E with the same experiment but continuous dilution. **B)** Fitness over time for selected *cbb* genes of the pHG1 encoded operon, except *cbbR* which is located on chromosome 2. Other chromosome 2 encoded *cbb* genes are not shown due to low transposon insertion frequency. Grayscale labels indicate role in CBB pathway: dark gray - transcriptional regulator, moderate gray - specific for $CO_2$ fixation, light gray - overlapping role in glycolysis/CBB cycle. Compare also Figure 5 F with the same experiment but continuous dilution. **C)** Fitness for all central carbon metabolism genes associated with the reactions in Figure 3 D. Genes are broken down by pathway. Dotted line - fitness ≤ -3 was regarded as significant. Compare also Figure 6 A with the same experiment but continuous dilution.

**Table S1. Summary of constraints for RBA model.**

| Parameter | Value | Unit | Reference |
|---|---|---|---|
| Biomass composition | adapted from genome scale model | g $gDCW^{-1}$ | Park et al., 2011 |
| replication efficiency (DNA pol III) | $2.88 \times 10^6$ | nt/h | average of multiple sources, see Methods |
| transcription efficiency (RNA pol II) | 223,200 | nt/h | Epshtein et al., 2003 |
| ribosome efficiency | 97,200 | aa/h | Bulovic et al., 2019 |
| chaperone efficiency | 36045 × μ - 2888 | aa/h | Bulovic et al., 2019 |
| growth-associated maintenance | 150 | mmol $gDCW^{-1}$ | Park et al., 2011 |
| non growth-associated maintenance | 3.0 | mmol $gDCW^{-1}$ $h^{-1}$ | Park et al., 2011 |
| total protein concentration (in aa) | 6.18 | mmol $gDCW^{-1}$ | Park et al., 2011 |
| fraction of cytoplasmic proteins | 0.8684 + 0.1060 × μ | unitless | this study |
| fraction of membrane proteins | 0.1316 - 0.1060 × μ | unitless | this study |
| fraction of non-enzymatic proteins in cytoplasm | 0.5374 - 0.5657 × μ | unitless | this study |
| fraction of non-enzymatic proteins in membrane | 0.8414 - 0.2147 × μ | unitless | this study |
| median $k_{app}$ | 5770 | $h^{-1}$ | this study |

**Table S2. Conditionally essential genes.** This table lists all annotated genes for the marked reactions in Figure 6 B, C, and D. Genes that were found to be essential in one or more conditions are marked with grey background. Fitness values below or above a threshold of |F| ≥ 3 are marked with red. Non-essential genes annotated for the same reactions are included for comparison.

| Pathway | Reaction | Reaction name | Gene ID | Gene name | for-mate | fruc-tose | succi-nate |
|---|---|---|---|---|---|---|---|
| CBB cycle | FDH | Formate dehydrogenase | H16_A0640 | fdsG | -4.1 | -0.1 | -0.1 |
| CBB cycle | FDH | Formate dehydrogenase | H16_A0641 | fdsB | -4.4 | 0.1 | 0 |
| CBB cycle | FDH | Formate dehydrogenase | H16_A0642 | fdsA | -5.3 | -0.2 | -0.1 |
| CBB cycle | FDH | Formate dehydrogenase | H16_A0644 | fdsD | -6.3 | 0.3 | 0.1 |
| CBB cycle | FDH | Formate dehydrogenase | H16_A2934 | fdhC | -0.4 | 0.3 | 0.4 |
| CBB cycle | FDH | Formate dehydrogenase | H16_A2936 | fdhB1 | -0.1 | 0 | 0.4 |
| CBB cycle | FDH | Formate dehydrogenase | H16_A2937 | fdhA1 | -0.2 | 0.1 | 0.1 |
| CBB cycle | FDH | Formate dehydrogenase | H16_A3292 | | -0.1 | -0.3 | -0.3 |
| CBB cycle | FDH | Formate dehydrogenase | H16_B1383 | cbbB | 0 | -0.3 | 0.3 |
| CBB cycle | FDH | Formate dehydrogenase | H16_B1452 | fdoG | -2.4 | 0.6 | 0.1 |
| CBB cycle | FDH | Formate dehydrogenase | H16_B1471 | fdhA2 | 0 | 0.1 | 0 |
| CBB cycle | FDH | Formate dehydrogenase | H16_B1700 | fdwA | -0.4 | 0.3 | -0.7 |
| CBB cycle | FDH | Formate dehydrogenase | H16_B1701 | fdwB | 0.2 | -0.6 | -0.7 |
| ED pathway | EDA | 2-dehydro-3-deoxy-phospho gluconate aldolase | H16_B1213 | eda | -2.6 | -4.3 | -1.7 |
| ED pathway | EDD | 6-phosphogluconate dehydratase | H16_A1178 | edd1 | -0.3 | -4.3 | -0.1 |
| ED pathway | EDD | 6-phosphogluconate dehydratase | H16_B2567 | edd2 | 0.1 | 0 | 0.2 |
| ED pathway | PGL | 6-phosphogluconolactonase | H16_B2565 | pgl | -2.2 | -3.9 | -1.1 |
| Pyruvate metabolism | ME1 | Malic enzyme (NAD) | H16_A3153 | maeA | 0 | 0.7 | -3.4 |
| Pyruvate metabolism | ME2 | Malic enzyme (NADP) | H16_A1002 | maeB | 0.3 | -0.2 | 0.2 |
| Pyruvate metabolism | PDH1 | Pyruvate dehydrogenase E1 component | H16_A1374 | pdhA | -2.6 | -5.2 | -3.2 |

| | | | | | | | |
|---|---|---|---|---|---|---|---|
| Pyruvate metabolism | PDH1 | Pyruvate dehydrogenase E1 component | H16_A1753 | pdhA2 | -0.4 | -1.1 | 0.1 |
| Pyruvate metabolism | PDH1 | Pyruvate dehydrogenase E1 component | H16_B0145 | acoB | -0.2 | -0.2 | 0.2 |
| Pyruvate metabolism | PDH1 | Pyruvate dehydrogenase E1 component | H16_B1300 | aceE | -0.1 | -0.3 | 0.4 |
| Pyruvate metabolism | PDH1 | Pyruvate dehydrogenase E1 component | H16_B2233 | bkdA1 | 0.2 | 0.2 | 0 |
| Pyruvate metabolism | PDH1 | Pyruvate dehydrogenase E1 component | H16_B2234 | bkdA2 | 0.2 | 0.3 | 0 |
| Pyruvate metabolism | PPC | Phosphoenolpyruvate carboxylase | H16_A2921 | ppc | -4.4 | -2.7 | -0.1 |

**Table S3. Primers used in this study**

| Primer Name | Primer Sequence | Primer Function | Source |
|---|---|---|---|
| Short_Biotin_pHIMAR | (Biot)-CGCCCTGCAGGGATGTCCACGAG | Biotinylated forward primer for amplify Tn-specific sequences during TnSeq | This work |
| NC102 | GTGACTGGAGTTCAGACGTGTGCTCTTCCGATC | Illumina I7 specific reverse primer for use with Short_Biotin_pHIMAR for amplifying Tn-specific sequences during TnSeq | This work |
| Nspacer_barseq_pHIMAR | ATGATACGGCGACCACCGAGATCTACACTCTTTCCCTACACGACGCTCTTCCGATCTNNNNNNCGCCCTGCAGGGATGTCCACGAG | Tn specific primer containing Illumina adaptors I5 and P5 as primer extensions. | Wetmore et al., 2015 |
| NEBNext Index 3 Primer for Illumina | CAAGCAGAAGACGGCATACGAGATGCCTAAGTGACTGGAGTTCAGACGTGTGCTCTTCCGATC- | Illumina I7 specific reverse primer containing an index sequence and an I7 adaptor. | New England Biolabs |
| BarSeq_R_P2_UMI_Univ | AATGATACGGCGACCACCGAGATCTACACTCTTTCCCTACACGACGCTCTTCCGATCTNNNNNNGTCGACCTGCAGCGTACG | Barcode specific reverse phasing primer 1, used as a pool with the other phasing primers during BarSeq. Adds Illumina I5 and P5 adaptors. | This work |
| BarSeq_R_P2_UMI_Univ_N2 | AATGATACGGCGACCACCGAGATCTACACTCTTTCCCTACACGACGCTCTTCCGATCTNNGTCGACCTGCAGCGTACG | Barcode specific reverse phasing primer 2, used as a pool with the other phasing primers during BarSeq. Adds Illumina I5 and P5 adaptors. | This work |
| BarSeq_R_P2_UMI_Univ_N3 | AATGATACGGCGACCACCGAGATCTACACTCTTTCCCTACACGACGCTCTTCCGATCTNNNGTCGACCTGCAGCGTACG | Barcode specific reverse phasing primer 3, used as a pool with the other phasing primers using BarSeq. Adds Illumina I5 and P5 adaptors. | This work |
| BarSeq_R_P2_UMI_Univ_N4 | AATGATACGGCGACCACCGAGATCTACACTCTTTCCCTACACGACGCTCTTCCGATCTNNNNGTCGACCTGCAGCGTACG | Barcode specific reverse phasing primer 4, used as a pool with the other phasing primers during BarSeq. Adds Illumina I5 and P5 adaptors. | This work |
| BarSeq_R_P2_UMI_Univ_N5 | AATGATACGGCGACCACCGAGATCTACACTCTTTCCCTACACG | Barcode specific reverse phasing primer 5, used as a | This work |

| | | | |
|---|---|---|---|
| | ACGCTCTTCCGATCTNNNNNG TCGACCTGCAGCGTACG | pool with the other phasing primers during BarSeq. Adds Illumina I5 and P5 adaptors. | |
| BarSeq_F_i7_001 | CAAGCAGAAGACGGCATACGA GATCGTGATGTGACTGGAGTT CAGACGTGTGCTCTTCCGATC TGATGTCCACGAGGTCTCT | Barcode specific forward indexing primer, used during BarSeq. Adds Illumina I7 and P7 adaptors. | This work |
| BarSeq_F_i7_002 | CAAGCAGAAGACGGCATACGA GATACATCGGTGACTGGAGTT CAGACGTGTGCTCTTCCGATC TGATGTCCACGAGGTCTCT | Barcode specific forward indexing primer, used during BarSeq. Adds Illumina I7 and P7 adaptors. | This work |
| BarSeq_F_i7_003 | CAAGCAGAAGACGGCATACGA GATGCCTAAGTGA CTGGAGTTCAGACGTGTGCTC TTCCGATCTGATGT CCACGAGGTCTCT | Barcode specific forward indexing primer, used during BarSeq. Adds Illumina I7 and P7 adaptors. | This work |
| BarSeq_F_i7_004 | CAAGCAGAAGACGGCATACGA GATTGGTCAGTGA CTGGAGTTCAGACGTGTGCTC TTCCGATCTGATGT CCACGAGGTCTCT | Barcode specific forward indexing primer, used during BarSeq. Adds Illumina I7 and P7 adaptors. | This work |
| BarSeq_F_i7_005 | CAAGCAGAAGACGGCATACGA GATCACTGTGTGA CTGGAGTTCAGACGTGTGCTC TTCCGATCTGATGT CCACGAGGTCTCT | Barcode specific forward indexing primer, used during BarSeq. Adds Illumina I7 and P7 adaptors. | This work |
| BarSeq_F_i7_006 | CAAGCAGAAGACGGCATACGA GATATTGGCGTGAC TGGAGTTCAGACGTGTGCTCT TCCGATCTGATGTC CACGAGGTCTCT | Barcode specific forward indexing primer, used during BarSeq. Adds Illumina I7 and P7 adaptors. | This work |
| BarSeq_F_i7_007 | CAAGCAGAAGACGGCATACGA GATGATCTGGTGA CTGGAGTTCAGACGTGTGCTC TTCCGATCTGATGT CCACGAGGTCTCT | Barcode specific forward indexing primer, used during BarSeq. Adds Illumina I7 and P7 adaptors. | This work |
| BarSeq_F_i7_008 | CAAGCAGAAGACGGCATACGA GATTCAAGTGTGAC TGGAGTTCAGACGTGTGCTCT TCCGATCTGATGTC CACGAGGTCTCT | Barcode specific forward indexing primer, used during BarSeq. Adds Illumina I7 and P7 adaptors. | This work |
| BarSeq_F_i7_009 | CAAGCAGAAGACGGCATACGA GATCTGATCGTGAC TGGAGTTCAGACGTGTGCTCT | Barcode specific forward indexing primer, used during BarSeq. Adds Illumina I7 and | This work |

| | | | |
|---|---|---|---|
| | TCCGATCTGATGTC CACGAGGTCTCT | P7 adaptors. | |
| BarSeq_F_i7_0 10 | CAAGCAGAAGACGGCATACGA GATAAGCTAGTGAC TGGAGTTCAGACGTGTGCTCT TCCGATCTGATGTC CACGAGGTCTCT | Barcode specific forward indexing primer, used during BarSeq. Adds Illumina I7 and P7 adaptors. | This work |
| BarSeq_F_i7_0 11 | CAAGCAGAAGACGGCATACGA GATGTAGCCGTGA CTGGAGTTCAGACGTGTGCTC TTCCGATCTGATGT CCACGAGGTCTCT | Barcode specific forward indexing primer, used during BarSeq. Adds Illumina I7 and P7 adaptors. | This work |
| BarSeq_F_i7_0 12 | CAAGCAGAAGACGGCATACGA GATTACAAGGTGAC TGGAGTTCAGACGTGTGCTCT TCCGATCTGATGTC CACGAGGTCTCT | Barcode specific forward indexing primer, used during BarSeq. Adds Illumina I7 and P7 adaptors. | This work |
| BarSeq_F_i7_0 13 | CAAGCAGAAGACGGCATACGA GATTTGACTGTGAC TGGAGTTCAGACGTGTGCTCT TCCGATCTGATGTC CACGAGGTCTCT | Barcode specific forward indexing primer, used during BarSeq. Adds Illumina I7 and P7 adaptors. | This work |
| BarSeq_F_i7_0 14 | CAAGCAGAAGACGGCATACGA GATGGAACTGTGA CTGGAGTTCAGACGTGTGCTC TTCCGATCTGATGT CCACGAGGTCTCT | Barcode specific forward indexing primer, used during BarSeq. Adds Illumina I7 and P7 adaptors. | This work |
| BarSeq_F_i7_0 15 | CAAGCAGAAGACGGCATACGA GATTGACATGTGAC TGGAGTTCAGACGTGTGCTCT TCCGATCTGATGTC CACGAGGTCTCT | Barcode specific forward indexing primer, used during BarSeq. Adds Illumina I7 and P7 adaptors. | This work |
| BarSeq_F_i7_0 16 | CAAGCAGAAGACGGCATACGA GATGGACGGGTGA CTGGAGTTCAGACGTGTGCTC TTCCGATCTGATGT CCACGAGGTCTCT | Barcode specific forward indexing primer, used during BarSeq. Adds Illumina I7 and P7 adaptors. | This work |
| BarSeq_F_i7_0 17 | CAAGCAGAAGACGGCATACGA GATCTCTACGTGAC TGGAGTTCAGACGTGTGCTCT TCCGATCTGATGTC CACGAGGTCTCT | Barcode specific forward indexing primer, used during BarSeq. Adds Illumina I7 and P7 adaptors. | This work |
| BarSeq_F_i7_0 18 | CAAGCAGAAGACGGCATACGA GATGCGGACGTGA | Barcode specific forward indexing primer, used during | This work |

| | | | |
|---|---|---|---|
| | CTGGAGTTCAGACGTGTGCTC TTCCGATCTGATGT CCACGAGGTCTCT | BarSeq. Adds Illumina I7 and P7 adaptors. | |
| BarSeq_F_i7_0 19 | CAAGCAGAAGACGGCATACGA GATTTTCACGTGAC TGGAGTTCAGACGTGTGCTCT TCCGATCTGATGTC CACGAGGTCTCT | Barcode specific forward indexing primer, used during BarSeq. Adds Illumina I7 and P7 adaptors. | This work |
| BarSeq_F_i7_0 20 | CAAGCAGAAGACGGCATACGA GATGGCCACGTGA CTGGAGTTCAGACGTGTGCTC TTCCGATCTGATGT CCACGAGGTCTCT | Barcode specific forward indexing primer, used during BarSeq. Adds Illumina I7 and P7 adaptors. | This work |
| BarSeq_F_i7_0 21 | CAAGCAGAAGACGGCATACGA GATCGAAACGTGA CTGGAGTTCAGACGTGTGCTC TTCCGATCTGATGT CCACGAGGTCTCT | Barcode specific forward indexing primer, used during BarSeq. Adds Illumina I7 and P7 adaptors. | This work |
| BarSeq_F_i7_0 22 | CAAGCAGAAGACGGCATACGA GATCGTACGGTGA CTGGAGTTCAGACGTGTGCTC TTCCGATCTGATGT CCACGAGGTCTCT | Barcode specific forward indexing primer, used during BarSeq. Adds Illumina I7 and P7 adaptors. | This work |
| BarSeq_F_i7_0 23 | CAAGCAGAAGACGGCATACGA GATCCACTCGTGA CTGGAGTTCAGACGTGTGCTC TTCCGATCTGATGT CCACGAGGTCTCT | Barcode specific forward indexing primer, used during BarSeq. Adds Illumina I7 and P7 adaptors. | This work |
| BarSeq_F_i7_0 24 | CAAGCAGAAGACGGCATACGA GATGCTACCGTGA CTGGAGTTCAGACGTGTGCTC TTCCGATCTGATGT CCACGAGGTCTCT | Barcode specific forward indexing primer, used during BarSeq. Adds Illumina I7 and P7 adaptors. | This work |
| BarSeq_F_i7_0 25 | CAAGCAGAAGACGGCATACGA GATATCAGTGTGAC TGGAGTTCAGACGTGTGCTCT TCCGATCTGATGTC CACGAGGTCTCT | Barcode specific forward indexing primer, used during BarSeq. Adds Illumina I7 and P7 adaptors. | This work |
| BarSeq_F_i7_0 26 | CAAGCAGAAGACGGCATACGA GATGCTCATGTGAC TGGAGTTCAGACGTGTGCTCT TCCGATCTGATGTC CACGAGGTCTCT | Barcode specific forward indexing primer, used during BarSeq. Adds Illumina I7 and P7 adaptors. | This work |
| BarSeq_F_i7_0 | CAAGCAGAAGACGGCATACGA | Barcode specific forward | This work |

| | | | |
|---|---|---|---|
| 27 | GATAGGAATGTGAC TGGAGTTCAGACGTGTGCTCT TCCGATCTGATGTC CACGAGGTCTCT | indexing primer, used during BarSeq. Adds Illumina I7 and P7 adaptors. | |
| BarSeq_F_i7_0 28 | CAAGCAGAAGACGGCATACGA GATCTTTTGGTGAC TGGAGTTCAGACGTGTGCTCT TCCGATCTGATGTC CACGAGGTCTCT | Barcode specific forward indexing primer, used during BarSeq. Adds Illumina I7 and P7 adaptors. | This work |
| BarSeq_F_i7_0 29 | CAAGCAGAAGACGGCATACGA GATTAGTTGGTGAC TGGAGTTCAGACGTGTGCTCT TCCGATCTGATGTC CACGAGGTCTCT | Barcode specific forward indexing primer, used during BarSeq. Adds Illumina I7 and P7 adaptors. | This work |
| BarSeq_F_i7_0 30 | CAAGCAGAAGACGGCATACGA GATCCGGTGGTGA CTGGAGTTCAGACGTGTGCTC TTCCGATCTGATGT CCACGAGGTCTCT | Barcode specific forward indexing primer, used during BarSeq. Adds Illumina I7 and P7 adaptors. | This work |
| BarSeq_F_i7_0 31 | CAAGCAGAAGACGGCATACGA GATATCGTGGTGAC TGGAGTTCAGACGTGTGCTCT TCCGATCTGATGTC CACGAGGTCTCT | Barcode specific forward indexing primer, used during BarSeq. Adds Illumina I7 and P7 adaptors. | This work |
| BarSeq_F_i7_0 32 | CAAGCAGAAGACGGCATACGA GATTGAGTGGTGA CTGGAGTTCAGACGTGTGCTC TTCCGATCTGATGT CCACGAGGTCTCT | Barcode specific forward indexing primer, used during BarSeq. Adds Illumina I7 and P7 adaptors. | This work |
| BarSeq_F_i7_0 33 | CAAGCAGAAGACGGCATACGA GATCGCCTGGTGA CTGGAGTTCAGACGTGTGCTC TTCCGATCTGATGT CCACGAGGTCTCT | Barcode specific forward indexing primer, used during BarSeq. Adds Illumina I7 and P7 adaptors. | This work |
| BarSeq_F_i7_0 34 | CAAGCAGAAGACGGCATACGA GATGCCATGGTGA CTGGAGTTCAGACGTGTGCTC TTCCGATCTGATGT CCACGAGGTCTCT | Barcode specific forward indexing primer, used during BarSeq. Adds Illumina I7 and P7 adaptors. | This work |
| BarSeq_F_i7_0 35 | CAAGCAGAAGACGGCATACGA GATAAAATGGTGAC TGGAGTTCAGACGTGTGCTCT TCCGATCTGATGTC CACGAGGTCTCT | Barcode specific forward indexing primer, used during BarSeq. Adds Illumina I7 and P7 adaptors. | This work |

| | | | |
|---|---|---|---|
| BarSeq_F_i7_036 | CAAGCAGAAGACGGCATACGAGATTGTTGGGTGACTGGAGTTCAGACGTGTGCTCTTCCGATCTGATGTCCACGAGGTCTCT | Barcode specific forward indexing primer, used during BarSeq. Adds Illumina I7 and P7 adaptors. | This work |
